# Dissecting FOXA1 pioneering function by acute pharmacological degradation

**DOI:** 10.64898/2026.02.25.707765

**Authors:** Lauren M. Hargis, Paige A. Barta, Yuxiang Zhang, Rachel E. Hayward, Benjamin F. Cravatt, Michael A. Erb

**Affiliations:** Department of Chemistry, The Scripps Research Institute, La Jolla, California, USA

## Abstract

Pioneer factors control transcription by opening chromatin, but a lack of chemical tools has made it difficult to study pioneer activity with kinetic precision. We recently reported covalent chemical probes that remodel the genomic localization of FOXA1, a prototypical pioneer factor essential for the growth of many breast and prostate cancers. Here, we expand the chemical toolbox for FOXA1 by developing a dTAG-based system for small molecule-induced FOXA1 degradation. Coupling pharmacological perturbations to rapid measurements of chromatin structure and function, we find that FOXA1 exclusively initiates chromatin opening at its genomic binding sites. Interestingly, this unidirectional outcome on accessibility both activates and represses gene transcription depending on the chromatin environment surrounding the FOXA1-binding sites. These effects apply to both androgen receptor (AR) target genes and other cancer-relevant genes. Our findings thus uncover regulatory features that translate FOXA1 pioneering activity into both activation and repression of transcriptional programs critical for cancer growth.

## Introduction

Transcription factors (TFs) bind enhancers and recruit transcriptional co-regulators to control gene expression programs that are critical in both development and disease^1^. While most TFs bind to their cognate DNA sequences only within accessible regions of the genome, a select class of pioneer factors can bind DNA in closed regions of chromatin and initiate chromatin opening^2–5^. FOXA1, a prototypical pioneer, plays a central role in establishing both physiologic and pathophysiologic gene regulatory networks in liver, prostate, and breast tissues^1,6–11^. This makes it an indispensable regulator of organismal development, as well as a compelling drug target for diseases like breast and prostate cancer^7,12^.

In prostate cancer, FOXA1 is often highly expressed and required for tumor growth and survival^10,13,14^. It binds lineage-specific enhancers prior to androgen stimulation and facilitates recruitment of the androgen receptor (AR) to a large subset of its genomic targets^10,15^. By establishing chromatin accessibility at AR-bound and other regulatory elements, FOXA1 helps define the enhancer landscape in prostate cancer cells and supports lineage-specific transcriptional programs^10^. Furthermore, recurrent somatic *FOXA1* mutations are common in both primary and advanced prostate tumors, particularly within the Wing2 region of the Forkhead (FKHD) DNA-binding domain, and these mutations can alter the DNA motif specificity of FOXA1 and redirect its binding to neomorphic *cis*-regulatory elements^16–20^.

Classically, FOXA1 is described as a transcriptional activator that functions by opening chromatin at enhancers. However, some genetic models have suggested that FOXA1 can also suppress transcription by occluding competing factors or contributing to repressive chromatin states^20–23^. It has been difficult to confidently resolve the mechanisms by which FOXA1 controls gene-specific transcriptional programs, due to the indirect, secondary effects that occur after prolonged genetic perturbations (such as mRNA knockdown, CRISPR-mediated gene knockout, or exogenous overexpression)^24–26^. For example, as shown by the Cancer Dependency Map, prolonged FOXA1 loss in prostate cancer models typically impairs cell growth ^27–29^, which complicates the use of genetic methods to identify the immediate gene regulatory functions of this pioneer factor and the mechanisms that underly them.

We recently reported the activity-based protein profiling (ABPP)-guided discovery of WX-02-23, a chiral small-molecule ligand that covalently engages cysteine 258 of FOXA1 in a stereoselective and DNA-dependent manner^30^. Located within the Wing2 region of the FKHD domain, C258 is positioned nearby the FOXA1-DNA interface and is adjacent to the most common cancer-associated FOXA1 mutations. WX-02-23 engagement of C258 improves FOXA1 binding to suboptimal DNA motifs and rapidly alters the genomic localization of FOXA1 in prostate cancer cells. The product of this reorganization is enhanced FOXA1 occupancy at many new sites in the genome that harbor suboptimal motifs alongside reductions in FOXA1 at sites bearing high-affinity motifs^30^. WX-02-23-induced changes in FOXA1 localization are positively correlated with changes in DNA accessibility, consistent with a model where FOXA1 binding is principally associated with the opening of chromatin. While WX-02-23 made it possible to study the relationship between FOXA1 localization and chromatin accessibility with temporal precision, additional chemical methods to disrupt FOXA1 activity genome-wide would enable a more thorough investigation of the effects of this pioneer factor on gene transcription.

Here, we use the dTAG (degradation tag) system to enable acute pharmacological loss of FOXA1 in prostate cancer cells. This FOXA1-dTAG model system relies on the genetic fusion of FOXA1 to FKBP12^F36V^, a “bump-and-hole” tag that can be targeted by small-molecule PROTACs (Proteolysis-Targeting Chimeras) to elicit rapid and highly selective degradation^31–33^. In 22Rv1 prostate cancer cells, we couple FOXA1 degradation with kinetically resolved genomic measurements of chromatin structure and function to better understand the direct mechanisms by which FOXA1 functions *in situ*. We find that FOXA1 degradation leads to the global closing of chromatin at tens of thousands of sites across the genome, and we did not observe evidence of FOXA1-binding sites becoming more accessible after FOXA1 loss. This general chromatin closing effect, however, resulted in nearly equivalent numbers of upregulated and downregulated genes, indicating that FOXA1 is both a unidirectional regulator of chromatin accessibility and a bidirectional regulator of gene transcription. FOXA1-binding sites that promote gene activation show greater amounts of FOXA1 enrichment, chromatin accessibility, and H3K27 acetylation, indicating that the local chromatin environment differentiates the opposing gene regulatory functions of FOXA1. Lastly, using WX-02-23 in combination with a SMARCA2/4 ATPase inhibitor, we find that FOXA1 can open chromatin independently of the SWI/SNF complex, further highlighting the utility of acute pharmacological perturbations for dissecting pioneer TF mechanisms in living cells.

## Results

### Rapid FOXA1 degradation enabled by the dTAG system

In pursuit of a rapid pharmacological approach for uniformly disrupting FOXA1 function genome-wide, we selected the dTAG system for targeted protein degradation. We first used the Cancer Dependency Map (DepMap 24Q4) to prioritize a cell line for FOXA1-dTAG development. Consistent with prior work implicating FOXA1 in AR- and ER-driven transcriptional programs^10,15,16,34–36^, the DepMap identified FOXA1 as one of the strongest selective dependencies in both prostate and breast cancer models (Figure 1A). Among the prostate cancer cell lines dependent on FOXA1, we chose 22Rv1 cells for establishing the dTAG system, as it expressed high levels of FOXA1 and showed strong dependency on this protein (Figure 1B). We confirmed the requirement of FOXA1 for growth of 22Rv1 cells by CRISPR/Cas9-based competitive proliferation experiments^37^, which track the fitness of knockout cells relative to wild-type cells (Figure 1C).

**Figure 1:**
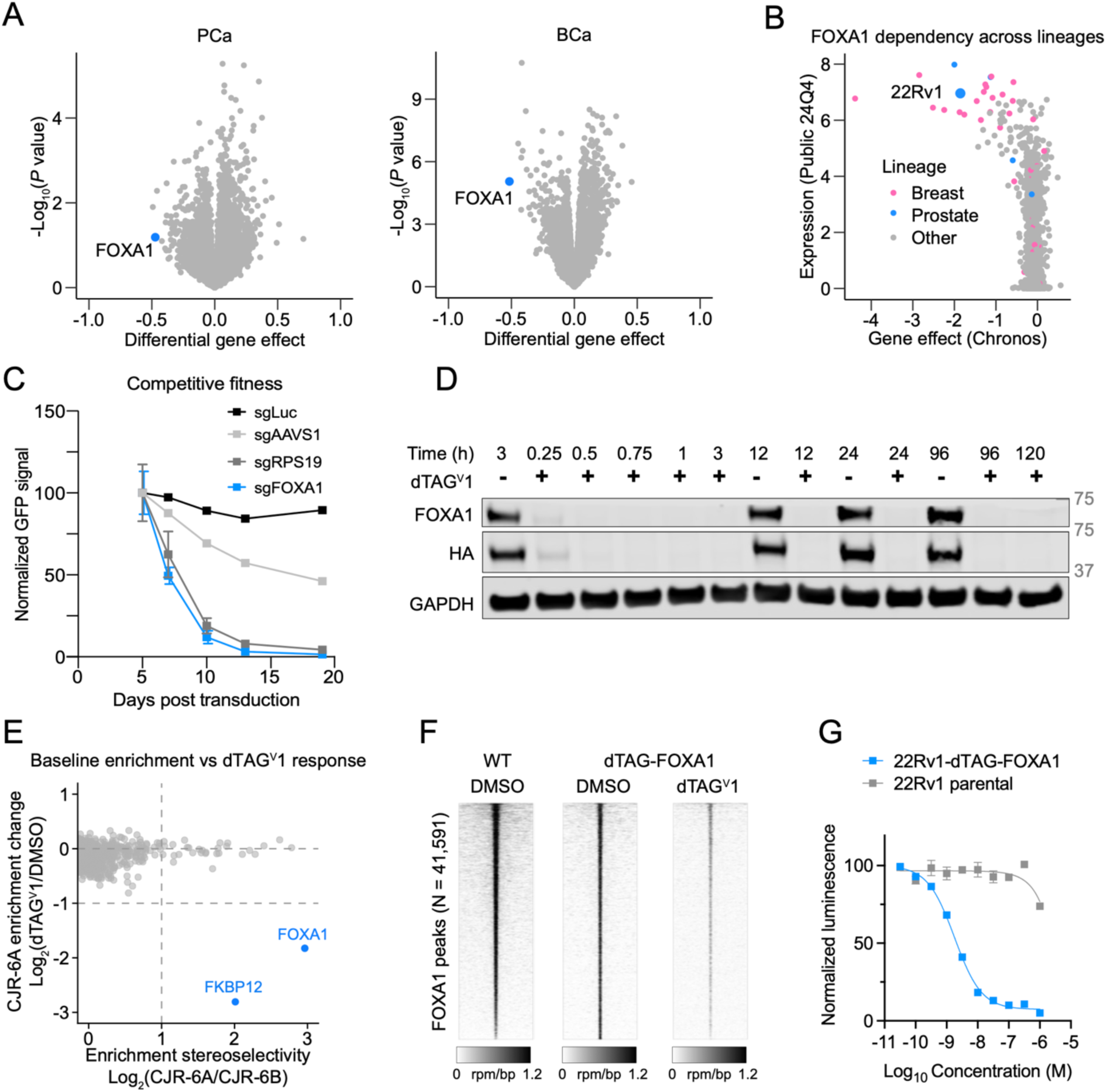
**A dTAG system for rapid FOXA1 degradation.** A. Volcano plots of differential gene dependencies in prostate and breast cancer cell. Left: differential dependencies were calculated for each gene using the average gene effect score (Chronos, DepMap, 24Q4) in prostate cancer (PCa) cell lines versus the average of all other cell lines. Right: breast cancer (BCa) cell lines versus the average of all other cell lines. B. FOXA1 gene expression (log-transformed TPM) versus gene effect score (Chronos) for each cell line in the DepMap dataset (24Q4). Each point represents an individual cell line. C. CRISPR-Cas9 competitive growth assays in 22Rv1-Cas9 cells. The proportion of GFP-positive cells was monitored by flow cytometry to track sgRNA-positive cells over time. Data represent mean ± SEM, *n* = 3 biological replicates. D. Time-course analysis of FOXA1 depletion. Immunoblot of FOXA1 and indicated proteins in 22Rv1-dTAG-FOXA1 cells treated with DMSO or 0.5 µM dTAG^V^1 for the indicated times. E. Scatter plot of protein-directed ABPP experiments showing stereoselective protein enrichment under DMSO conditions at 1 h (log₂[CJR-6A/CJR-6B]) and the change in enrichment by CJR-6A following dTAG^V^1 treatment (log₂[dTAG^V^1/DMSO]). Cells were treated with DMSO or dTAG^V^1 (500 nM, 0.5 h), followed by labeling with CJR-6A or CJR-6B (5 µM, 1 h). Data are shown as mean ± SEM (*n* = 2 independent experiments for CJR-6B; *n* = 3 independent experiments for all other conditions). F. Heatmaps displaying ChIP-seq signal intensity across combined FOXA1 (wild-type 22Rv1 cells, GSE261803)^30^ and dTAG-FOXA1 (22Rv1-dTAG-FOXA1 cells) ChIP-seq peaks. All samples are rank ordered by the wild-type FOXA1 sample. 22Rv1-dTAG-FOXA1 cells were treated for 1.5 h with 500 nM dTAG^V^1 or DMSO)^30^. Data are representative of 2 independent experiments. G. Effect of dTAG^V^1 on growth of wild-type 22Rv1 and 22Rv1-dTAG-FOXA1 cells (5-day treatment, CellTiter-Glo signal normalized to DMSO with subtraction of day 0 values). Data represent mean ± SEM, *n* = 3 biological replicates.

FKBP12^F36V^ and a tandem HA epitope tag were fused to the amino (N)-terminus of FOXA1 using CRISPR/Cas9 and homology-directed repair (Figure S1A). A mixture of homology donor templates encoding resistance to blasticidin and puromycin was introduced (Table S1), and cells with biallelic integrations were enriched by sequential antibiotic selection (Figure S1A). The wild-type *FOXA1* allele was not detectable by PCR, nor was its expression observed by immunoblot analysis, confirming successful selection of a population of cells with biallelic knock-ins (Figures S1B and S1C). In these cells, treatment with dTAG^V^1, a PROTAC designed from ligands for FKBP12^F36V^ and the E3 substrate receptor VHL (Von Hippel-Lindau)^38^, induced rapid and concentration-dependent degradation of FOXA1 (Figures 1D and S1C). Maximal degradation occurred within 30-45 min and was sustained for at least 5 days after a single exposure to dTAG^V^1 (Figure 1D), thus establishing a system for temporally precise and durable perturbations of FOXA1.

Protein-directed ABPP revealed stereoselective engagement of dTAG-FOXA1 by CJR-6A^30^, an alkyne-functionalized analog of WX-02-23, thus providing initial evidence that the dTAG-FOXA1 fusion protein was properly folded in 22Rv1 cells (Figure S1D, Supplementary Dataset S1). Treatment with dTAG^V^1 for 1.5 h markedly reduced the enrichment of FOXA1 and FKBP12 in protein-directed ABPP experiments, while other proteins enriched by CJR-6A were unperturbed (Figure 1E, Supplementary Dataset S1), supporting the selective targeted degradation of FOXA1. Similarly, ChIP-seq analyses showed that dTAG-FOXA1 localizes to endogenous FOXA1 binding sites in 22Rv1 cells and its binding to the genome was dramatically reduced by dTAG^V^1 treatment (Figures 1F and S1E-F, Supplementary Dataset S2). Although low residual signal remained detectable by ChIP-seq, likely reflecting the high sensitivity of enrichment-based assays, global FOXA1 binding was substantially diminished. These data supported that dTAG-FOXA1 is functional in 22Rv1 prostate cancer cells, which we further confirmed in cell proliferation assays, where dTAG^V^1 impaired the growth of 22Rv1-dTAG-FOXA1 cells, but not parental 22Rv1 cells (Figures 1G and S1G).

### FOXA1 binding promotes chromatin accessibility

To investigate the relationship between FOXA1 binding and chromatin accessibility on a global scale, we performed ATAC-seq on 22Rv1-dTAG-FOXA1 cells following acute treatment with dTAG^V^1 (1.5 h). Of the 40,627 peaks detected by ATAC-seq, more than 3,000 sites exhibited significant decreases in accessibility following dTAG^V^1 treatment (Log_2_FC < −1, *P* < 0.05), while fewer than 700 sites showed increases in accessibility (Log_2_FC > 1, *P* < 0.05) (Figure 2A, Supplementary Dataset S2). Stratification of ATAC-seq peaks by FOXA1 binding status using the dTAG-FOXA1 ChIP-seq dataset revealed significantly greater losses in chromatin accessibility at FOXA1-bound sites compared to unbound sites (Figure 2B, Supplementary Dataset S2). Nearly 98% of ATAC-seq decreases overlapped with d-TAG-FOXA1 ChIP-seq peaks, whereas less than 2% of accessibility increases did so (Figure 2C, Supplementary Dataset S2), indicating that chromatin closing represents the primary direct consequence of FOXA1 degradation, while accessibility gains are likely indirect. Indeed, focusing on the ATAC-seq signals at all 32,322 FOXA1-binding sites, we found that decreases after dTAG^V^1 treatment showed a strong positive correlation with the degree of dTAG-FOXA1 depletion from these sites (Figures S2A and S2B, Supplementary Dataset S2), further indicating that the chromatin accessibility changes are directly caused by FOXA1 degradation.

**Figure 2:**
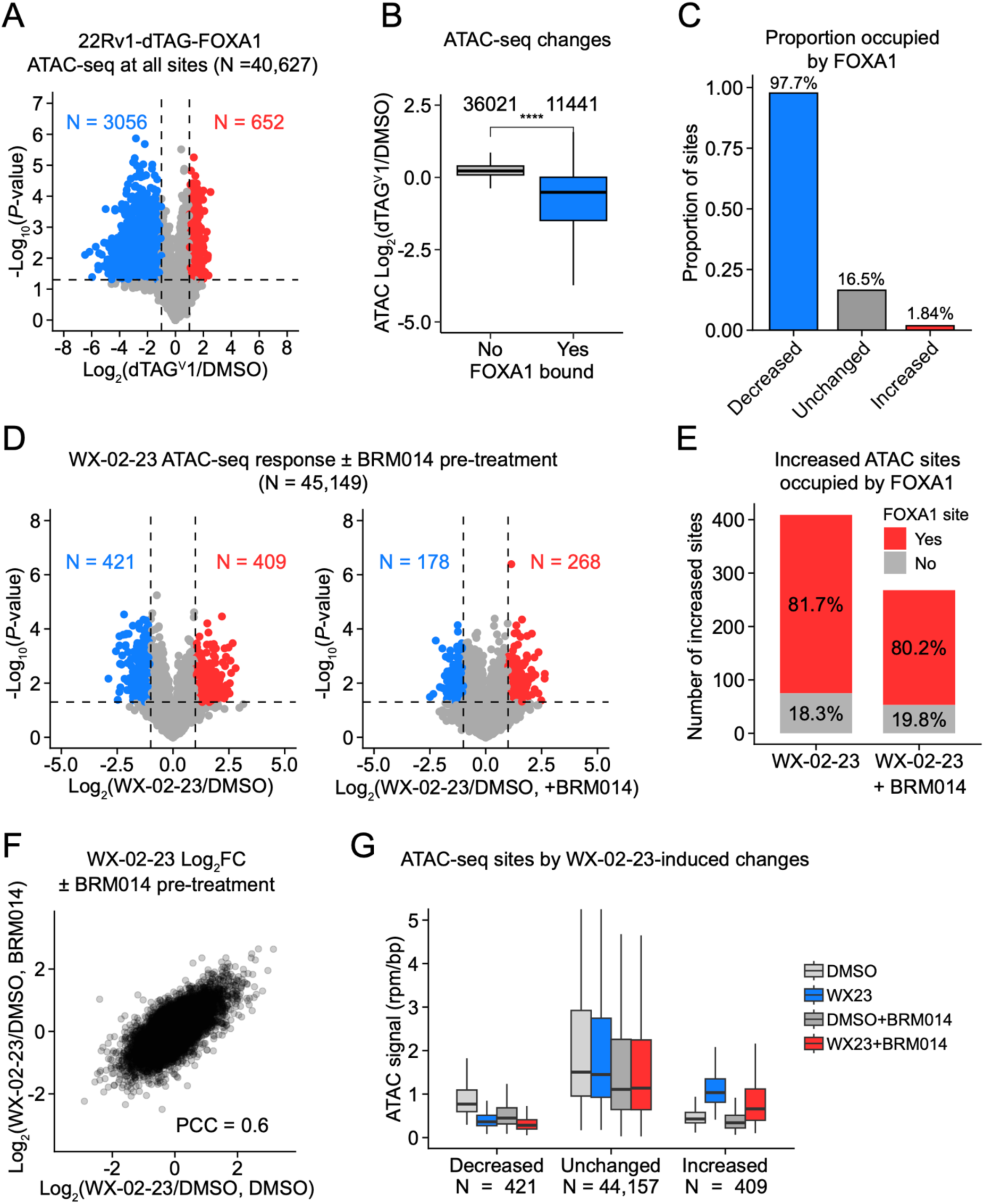
**Acute FOXA1 degradation uncovers a unidirectional effect on chromatin accessibility.** A. Chromatin accessibility (ATAC-seq) changes following dTAG^V^1 treatment (DMSO or 500 nM dTAG^V^1, 1 h) at all ATAC-seq sites (*N* = 40,627) in 22Rv1-dTAG-FOXA1 cells. ATAC-seq peaks with substantial (|log₂FC| > 1) and significant (*P* < 0.05) changes are shown in blue (decreased) or red (increased). *P* values determined by two-tailed Welch’s *t*-tests. *n* = 3 biological replicates for ATAC-seq. B. Boxplots showing changes in chromatin accessibility in 22Rv1-dTAG-FOXA1 cells at ATAC-seq sites with and without FOXA1 binding (as determined by overlap with dTAG-FOXA1 ChIP-seq peaks). Numbers above boxes indicate the number of sites per group. *n* = 3 biological replicates for ATAC-seq. Data are representative of 2 independent experiments. C. Proportion of ATAC-seq sites that overlap with dTAG-FOXA1 ChIP-seq peaks (in 22Rv1-dTAG-FOXA1). Sites are grouped by ATAC-seq responses to dTAG^V^1 (decreased: log_2_FC < −1, *P* < 0.05; unchanged: |log_2_FC| < 1 and *P* > 0.05; increased: log_2_FC > 1, *P* < 0.05). D. Volcano plots of ATAC-seq changes in 22Rv1-SF3B1-C1111S cells. Left: 20 µM WX-02-23 without BRM014 (1-h DMSO pretreatment, followed by WX-02-23 for 3 h). Right: 20 µM WX-02-23 following BRM014 pretreatment (1 µM BRM014 for 1 h, followed by WX-02-23 for 3 h). Significantly changed sites (|log₂FC| > 1, *P* < 0.05) are shown in blue (decreased) or red (increased), with counts labeled. *n* = 3 biological replicates for ATAC-seq. E. Overlap between ATAC-seq sites significantly increased by WX-02-23 (with or without BRM014, defined in 4D) and FOXA1 ChIP-seq peaks (parental 22Rv1 cells treated with DMSO or WX-02-23, GSE261803)^30^. F. Scatter plot comparing ATAC-seq following WX-02-23 treatment with or without 1 µM BRM014 pretreatment at all ATAC-seq sites (1-h DMSO or BRM014 followed by 3-h WX-02-23). PCC: Pearson correlation coefficient. *n* = 3 biological replicates for ATAC-seq. G. Boxplot of ATAC-seq signals (rpm/bp) at sites categorized by their response to WX-02-23 without BRM014 pretreatment: significantly decreased (log₂FC < −1 and *P* < 0.05, *N* = 421), unchanged (|log₂FC| < 1 and *P* > 0.05, *N* = 44,157), or significantly increased (log₂FC > 1 and *P* < 0.05, *N* = 409). Boxes represent median and interquartile range (IQR); whiskers indicate 1.5 × IQR. *n* = 3 biological replicates for ATAC-seq.

Biochemical studies have demonstrated that FOXA1 can open chromatin locally without the assistance of ATP-dependent chromatin remodelers *in vitro*^3,5,11,39^. While this is thought to occur in cells as well, the relative contribution of pioneer factors and chromatin remodelers to chromatin accessibility can be difficult to deconvolute. Our previous FOXA1 IP-MS experiments revealed a pronounced enrichment of SWI/SNF subunits in 22Rv1 cells^30^, consistent with prior work showing collaboration between pioneers and chromatin remodelers to maintain stable chromatin accessibility after an initial seeding event^40,41^. Evidence supporting an important role for chromatin remodelers include the global suppression of enhancer accessibility caused by pharmacological inhibition of the SWI/SNF complex^42–44^, and the disruption of FOXA1 engagement at its genomic binding sites in cells treated with SMARCA2/4 degraders^42^. To help deconvolute the contributions of FOXA1 and SWI/SNF to chromatin accessibility, we sought to determine if FOXA1 binding sites created by WX-02-23 could be opened in the absence of SWI/SNF activity.

For this experiment, we used 22Rv1 cells expressing a C1111S mutant form of the splicing factor SF3B1 that engenders resistance to reactivity with WX-02-23 and the corresponding perturbations in splicing observed in cells expressing WT-SF3B1^30,45^. C1111S-SF3B1 cells were treated with the SMARCA2/4 ATPase inhibitor BRM014 for 1 h prior to treatment with DMSO or WX-02-23 for an additional 3 h. As expected, SMARCA2/4 ATPase inhibition produced a widespread decrease in chromatin accessibility (Figure S2C, Supplementary Dataset S2). WX-02-23, however, still altered chromatin accessibility in cells treated with BRM014, albeit to a less dramatic extent compared to control cells (Figures 2D, Supplementary Dataset S2). In both the presence and absence of BRM014, more than 80% of the WX-02-23–induced increases in chromatin accessibility overlap with FOXA1 binding sites (as defined by previous ChIP-seq experiments of FOXA1 in DMSO and WX-02-23-treated 22Rv1 cells) (Figure 2E, Supplementary Dataset S2). Thus, WX-02-23-induced increases in chromatin accessibility remain direct effects of FOXA1 binding even in the presence of BRM014, as further supported by the strong correlation between ATAC-seq responses to WX-02-23 in the presence and absence of BRM014 (Figure 2F, Supplementary Dataset S2). After grouping ATAC-seq sites that are increased, decreased, or unchanged by WX-02-23 in the absence of BRM014, we found that WX-02-23 caused similar ATAC-seq changes to these groups in the presence of BRM014, despite the uniformly depressed levels of chromatin accessibility caused by BRM014 (Figures 2G, Supplementary Dataset S2). These findings demonstrate that both FOXA1 and SWI/SNF contribute to maximizing chromatin accessibility, but FOXA1 can also independently increase local chromatin accessibility even when SWI/SNF ATPase activity is inhibited.

### FOXA1 chromatin opening supports transcriptional activation and repression

We performed 3’-end mRNA-seq in 22Rv1-dTAG-FOXA1 cells following treatment with dTAG^V^1 for 3, 8, or 24 h to determine the transcriptional consequences of losing FOXA1 pioneering function (Figure 3A, Supplementary Dataset S2). In contrast to the unidirectional effects of FOXA1 degradation on chromatin accessibility, the ensuing transcriptional changes were bidirectional, resulting in relatively balanced numbers of up- and down-regulated genes across all time points (Figure 3A, Supplementary Dataset S2). Consistent with a role for FOXA1 in supporting cancer cell growth and malignancy, gene set enrichment analysis revealed downregulation of proliferation- and epithelial to mesenchymal (EMT)-associated gene signatures and upregulation of tumor-suppressive pathways following loss of this pioneer factor (Figure 3B, Supplementary Dataset S2). Interestingly, these transcriptional changes accumulated gradually over time, with fewer than 300 genes significantly altered at early time points and the greatest number of differentially expressed transcripts observed at 24 h (Figure 3C, Supplementary Dataset S2). Notably, several prostate lineage-specific genes, including *HOXA9**/**10*, *KLK2*, *SOX9*, and *NKX3-1*, were downregulated across all timepoints (Figure 3D, Supplementary Dataset S2), indicating that FOXA1 contributes to maintenance of a prostate-specific gene regulatory network. Conversely, *PDK4, ADRA1A, IGFBP3, ETS2, MME,* and *CRYAB* were upregulated following FOXA1 degradation, consistent with activation of stress-response and growth-suppressive transcriptional programs (Figure 3D, Supplementary Dataset S2).

**Figure 3:**
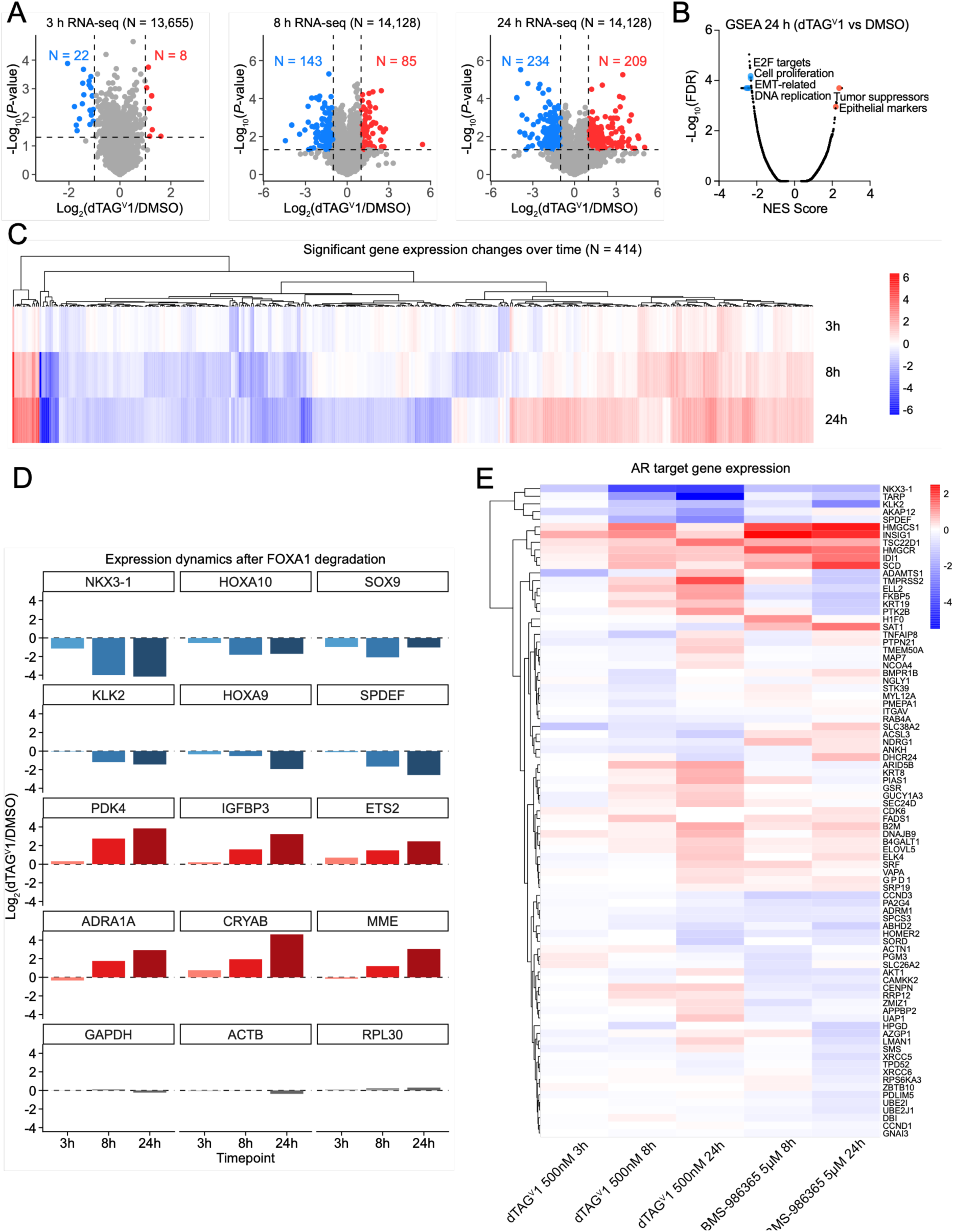
**FOXA1 degradation causes bi-directional changes in transcription.** A. mRNA-seq data showing gene expression changes in 22Rv1-dTAG-FOXA1 cells treated with 500 nM dTAG^V^1 or DMSO for 3, 8 or 24 h. Transcripts significantly altered by dTAG^V^1 were defined as |log₂(dTAG^V^1/DMSO)| > 1 and *P* < 0.05, with red or blue for increased or decreased, respectively. All transcripts with substantial signal (> 3 average CPM) in at least one treatment group were included (*N* = 13,655 for 3h, 14,128 transcripts for 8 and 24h). *n* = 3 biological replicates. *P* values were determined by two-tailed Welch’s *t*-test. B. Gene Set Enrichment Analysis (GSEA) of significantly altered genes (|log₂(dTAG^V^1/DMSO)| > 1, *P* < 0.05) after 24-h dTAG^V^1 treatment. Hallmark and curated (C2) gene sets were used from the molecular signatures database (MSigDB). Scatter plot of normalized enrichment score (NES) and false discovery rate (FDR) q-values. *n* = 3 biological replicates. C. Heatmap showing the combined list of significantly changed transcripts across all time points. Transcripts were hierarchically clustered and included if |log₂(dTAG^V^1/DMSO)| > 1 and *P* < 0.05 at any timepoint. *n* = 3 biological replicates. *P* values were calculated using two-tailed Welch’s *t*-test. D. Bar plots depicting transcript changes induced by dTAG^V^1 for selected prostate-associated genes. Housekeeping genes are represented in grey, downregulated genes in shades of blue, and upregulated genes in shades of red. E. Heatmap of transcript changes for AR target genes (GSE275777) following dTAG^V^1 or BMS-986365 treatment.

We found that the bidirectional impact of FOXA1 degradation extended to genes regulated by AR, which is known to functionally and physically interact with FOXA1^10,15^ (Figure 3E, Supplementary Dataset S2). Interestingly, the effects of FOXA1 degradation were, in several instances, counter to the known directionality of AR inhibition^46^. To confirm this, we evaluated the dual AR PROTAC and antagonist BMS-986365^47,48^ in 22Rv1-dTAG-FOXA1 cells (Figures 3E, S3A, and S3B, Supplementary Dataset S2). We found that several established AR target genes, including *FKBP5*, *TMPRSS2*, and *ELL2*, were downregulated by BMS-986365 while being upregulated by dTAG^V^1 (Figure S3C, Supplementary Dataset S2). Together, these findings demonstrate that FOXA1 and AR degradation yield both shared and distinct transcriptional outcomes, indicating that FOXA1 does not simply augment AR-dependent gene control but instead modulates AR targets in a context-dependent manner.

### Context-dependent enhancer regulation by FOXA1

We next compared the features of FOXA1 binding sites associated with upregulated vs downregulated target genes. ROSE2 was used to stitch proximal FOXA1 ChIP-seq peaks and assign putative target genes based on proximity. Changes in FOXA1 occupancy or chromatin accessibility at FOXA1 binding sites did not strongly correlate with the direction of transcriptional responses at associated gene targets (Figures 4A and 4B, Supplementary Dataset S2), indicating that the distinct transcriptional outcomes among FOXA1 targets were not due to locus-specific differences in the degree of FOXA1 loss following dTAG^V^1-induced degradation. On the other hand, basal levels of FOXA1 occupancy and chromatin accessibility were found to be higher at FOXA1 binding sites linked to target genes that were downregulated upon FOXA1 degradation (Figure 4C, Supplementary Dataset S2). Despite these differences in basal chromatin accessibility at the FOXA1 binding sites associated with gene activation versus repression (Figure 4D), FOXA1 degradation decreased the accessibility of both groups equivalently (Figure 4B, Supplementary Dataset S2). This finding indicates that FOXA1 fulfills a similar role in opening chromatin at both types of FOXA1 regulatory elements, with other factors likely contributing to the overall differences in chromatin accessibility observed at these sites.

**Figure 4.**
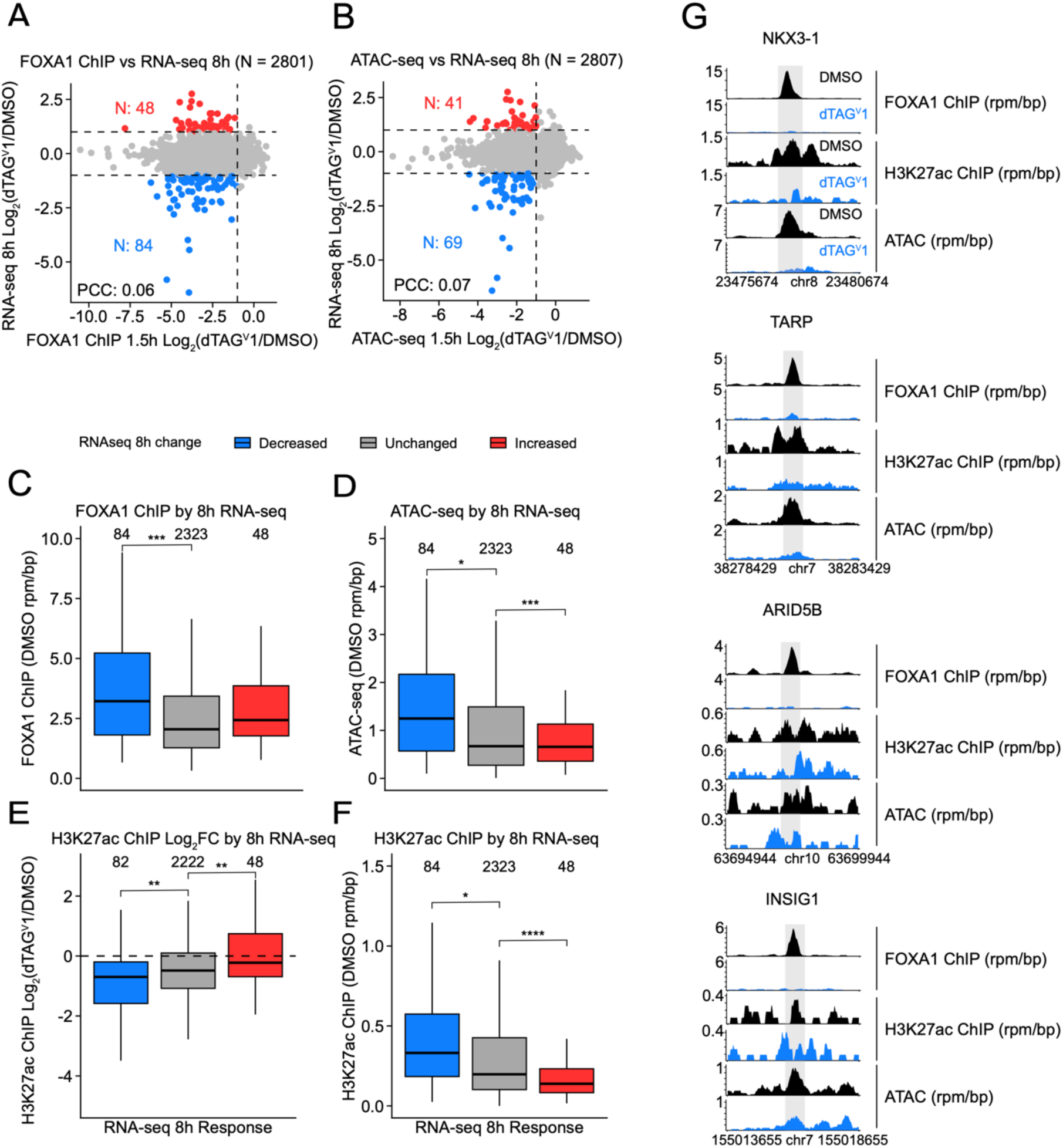
**Context-dependent transcriptional regulation by FOXA1 binding sites.** A. Scatter plot comparing changes in dTAG-FOXA1 ChIP-seq signal at dTAG-FOXA1 binding sites versus 8 h RNA-seq gene expression changes at their putative target genes (*N* = 2801 element-gene pairs). Element-gene pairs with 2-fold loss in dTAG-FOXA1 ChIP-seq signal and a 2-fold change in transcription are shown in red (increased mRNA expression) or blue (decreased mRNA expression). B. Scatter plot comparing changes in ATAC-seq signal at dTAG-FOXA1 binding sites versus 8 h RNA-seq gene expression changes at their putative target genes (*N* = 2807 element-gene pairs). C. dTAG-FOXA1 ChIP-seq signal (rpm/bp from DMSO-treated sample) at dTAG-FOXA1 binding sites. Box plots were grouped by the element-gene pairs showing 2-fold loss in dTAG-FOXA1 ChIP-seq signal and 2-fold increase (red) or decrease (blue) in mRNA-seq transcript abundance (dTAG^V^1, 8 h), as described in A. D. Same as C, but for ATAC-seq (rpm/bp from DMSO-treated samples) at dTAG-FOXA1 binding sites. Element-gene pairs were grouped as in A. E. Changes in H3K27ac ChIP-seq signal at dTAG-FOXA1 binding sites following dTAG^V^1 treatment (500 nM, 1.5 h), with element-gene pairs grouped as in A. F. Same as C, but for H3K27ac ChIP-seq signal (rpm/bp in DMSO-treated sample) at FOXA1 binding sites. Element-gene pairs were grouped as in A. G. Genome tracks showing FOXA1 binding sites linked to transcripts repressed (*NKX3-1, TARP*) or activated (*ARID5B, INSIG1*) by dTAG^V^1 treatment.

In considering the other chromatin features that might contribute to the differential transcriptional consequences of FOXA1 degradation, we used ChIP-seq to measure the abundance of H3K27ac, a mark of active enhancers^49^, with and without FOXA1 degradation. H3K27ac ChIP-seq profiles showed that global acetylation levels remained broadly stable following FOXA1 degradation (Figures S4A and S4B, Supplementary Dataset S2). However, H3K27ac sites also bound by FOXA1 exhibited significant reductions in H3K27ac ChIP-seq signal following dTAG^V^1 treatment (Figure S4C, Supplementary Dataset S2). These data indicate that FOXA1 is important for the maintenance of active enhancers at a large fraction of its binding sites. Indeed, FOXA1 binding sites linked to gene activation showed the highest basal levels of H3K27ac and the corresponding greatest reductions in these signals following FOXA1 degradation (blue bars, Figure 4E and 4F, Supplementary Dataset S2). In contrast, the lowest levels of basal H3K27ac were seen at FOXA1 binding sites associated with gene repression (red bars, Figure 4F, Supplementary Dataset S2). For example, FOXA1 binding sites proximal to *NKX3-1* and *TARP*, which are both downregulated after FOXA1 loss, exhibited coordinated decreases in FOXA1 binding, chromatin accessibility, and H3K27ac (Figure 4G, Supplementary Dataset S2). In contrast to these sites, FOXA1 binding sites proximal to *ARID5B* and *INSIG1*, which are upregulated after the loss of FOXA1, displayed lower basal levels of chromatin accessibility and H3K27ac and more muted decreases following FOXA1 degradation (Figure 4G, Supplementary Dataset S2). Similarly, AR occupancy is highest at sites linked to genes that are activated by FOXA1 (Figure S4D, Supplementary Dataset S2). Together with the observation that FOXA1 loss upregulates a subset of AR target genes, these findings suggest that the influence of FOXA1 on AR output varies depending on local enhancer context, with stronger AR occupancy associated with transcriptional activation.

## Discussion

In this study, we combined rapid pharmacological degradation of FOXA1 with kinetically resolved genomic measurements of chromatin structure and function to uncover direct mechanisms of FOXA1-dependent gene control in prostate cancer cells^23^. Our findings provide direct evidence that FOXA1 loss leads to both activation and repression of its target genes, despite causing a consistent decrease in chromatin accessibility at its genomic binding sites. This supports a model in which FOXA1 contributes broadly to the accessibility of *cis*-regulatory elements but functions as either a transcriptional activator or repressor depending on the local chromatin environment or higher order regulatory mechanisms. The lack of chromatin accessibility increases following FOXA1 degradation refines prevailing models that FOXA1 may directly close chromatin at repressive elements^3,21,23^, instead supporting a model in which FOXA1 binding sites exist in a partially open or poised state that requires continuous FOXA1 occupancy to remain accessible while other regulatory factors help mediate downstream transcriptional outcomes.

Previous studies of FOXA1 function have largely relied on overexpression systems or chronic knockdown/knockout models, which can obscure immediate regulatory effects of this pioneer factor and introduce indirect transcriptional changes from cellular adaptation. Such approaches have yielded conflicting conclusions on whether FOXA1 primarily functions as an activator, a repressor, or both, and on the extent to which chromatin accessibility changes accompany transcriptional outcomes. By employing the dTAG system for rapid, near-complete FOXA1 degradation, we can minimize confounding adaptive responses and reveal the acute, primary consequences of FOXA1 loss. Within 30 minutes of FOXA1 degradation, chromatin accessibility is decreased at nearly all FOXA1-binding sites. These rapid changes indicate that FOXA1 is not merely an initiating pioneer factor but also continuously required to maintain chromatin accessibility^11^.

Our data further clarify the relationship between FOXA1 and SWI/SNF chromatin remodeling complexes. While basal accessibility at some FOXA1-bound enhancers requires BRG1/BRM ATPase activity, most accessibility changes induced by the FOXA1 ligand WX-02-23 persist in the presence of the SMARCA2/4 ATPase inhibitor, BRM014. This suggests that FOXA1 can maintain chromatin accessibility through mechanisms independent of SWI/SNF catalytic activity, potentially involving other remodelers or local nucleosome destabilization via its winged-helix domain. This finding adds to emerging evidence that pioneer factors may stabilize open chromatin states through multiple, partially redundant mechanisms, and that their influence is not uniformly SWI/SNF-dependent^11,50–52^.

A striking outcome of our integrative analysis is that loss of chromatin accessibility caused by FOXA1 depletion does not dictate a single transcriptional directionality. FOXA1 removal leads to both activation and repression of its target genes, even though both classes reflect decreases in chromatin accessibility. This context-dependent transcriptional outcome suggests that FOXA1 can act as either an activator or a repressor at different *cis*-regulatory elements, with the direction presumably determined by other mechanisms, such as the presence of other co-regulatory factors, local chromatin features, and genome topology. For repressed genes, FOXA1 may stabilize a partially open chromatin state that is nonetheless incompatible with transcriptional activation, with its removal relieving this constraint. In contrast, for activated genes, FOXA1 loss is typically accompanied by reduced H3K27ac and accessibility at associated enhancers, supporting a direct role in maintaining active regulatory environments at these sites.

In sum, by capturing the immediate consequences of FOXA1 loss, our work advances our understanding of pioneer factor function and clarifies the role of FOXA1 in maintaining chromatin accessibility within lineage-specific transcriptional networks. The consistent downregulation of prostate lineage-defining genes such as *NKX3-1*, *TARP*, and *SOX9* in FOXA1-depleted cells underscores the essential role of this pioneer factor in sustaining prostate epithelial identity, which is reinforced by the concurrent upregulation of tumor-suppressive pathways. The observation that FOXA1 degradation alone impairs prostate cancer cell proliferation, together with the lineage-restricted essentiality of FOXA1 in DepMap data, supports the concept that targeting FOXA1 could selectively compromise prostate cancer pathogenesis, further motivating FOXA1 as a target for therapeutic discovery.

## Acknowledgements

This work was supported by the Ono Pharma Foundation Breakthrough Science Initiative Awards Program (M.A.E.), the Baxter Foundation Young Investigator Award (M.A.E.), the National Cancer Institute (R01CA280720, M.A.E.; R35CA231991, B.F.C.), and the David C. Fairchild Endowed Fellowship in the Skaggs Graduate School of Chemical and Biological Sciences (L.M.H.). We thank the Flow Cytometry Core at The Scripps Research Institute for assistance with cell sorting; the Genomics Core at The Scripps Research Institute, including John Shimashita, Sheila Roberts Wieland, Jessica Ledesma, and Steven Robert Head, for sequencing support; and Bruno Melillo for helpful discussions on data analysis.

---

**Figure S1:**
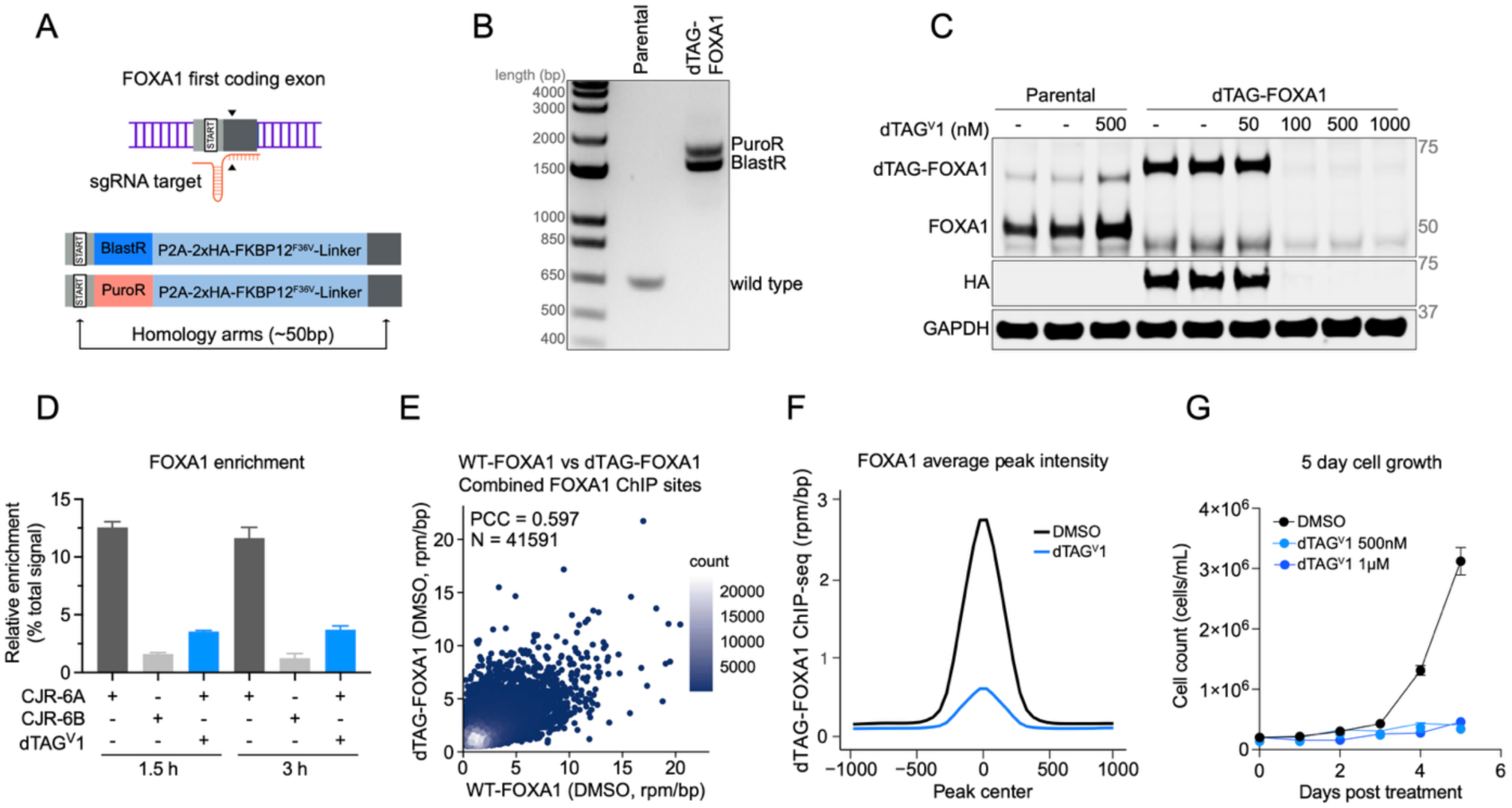
A dTAG system for rapid FOXA1 degradation, related to. **Figure 1**. A. Schematic of dTAG-FOXA1 engineering strategy. Illustration of the FOXA1 sgRNA target site at the N-terminal coding exon and the donor templates containing 2xHA-FKBP12^F36V^ degron cassettes with either blasticidin or puromycin resistance genes. Full donor sequences can be found in Table S1. B. PCR validation of FOXA1 degron integration. Gel electrophoresis showing PCR products from 22Rv1 wild-type or dTAG-FOXA1 knock-in cells (ethidium bromide stain). C. Immunoblot analysis of FOXA1 degradation showing FOXA1 protein levels in 22Rv1 wild-type and dTAG-FOXA1 cells treated with DMSO or dTAG^V^1 for 1 h. D. Protein-directed ABPP data for FOXA1 from 22Rv1-dTAG-FOXA1 cells treated with DMSO or dTAG^V^1 (500 nM, 0.5 or 2 h) followed by CJR-6A or CJR-6B (5 µM, 1 h). Average ± SEM (*n* = 2 independent experiments for CJR-6B, *n* = 3 independent experiments for all others). E. Scatter plot comparing WT-FOXA1 (GSE261803)^30^ vs dTAG-FOXA1 ChIP-seq signal (rpm/bp) across combined ChIP-seq peaks (*N* = 41,491). F. Averaged FOXA1 ChIP-seq signal (rpm/bp) at dTAG-FOXA1 ChIP-seq peaks in 22Rv1-dTAG-FOXA1 cells. G. Cell growth kinetics of 22Rv1-dTAG-FOXA1 cells treated with DMSO or dTAG^V^1 (500 nM or 1 µM). Cell counts were determined using a Countess automated cell counter and plotted as mean ± SD (*n* = 3 biological replicates).

**Figure S2:**
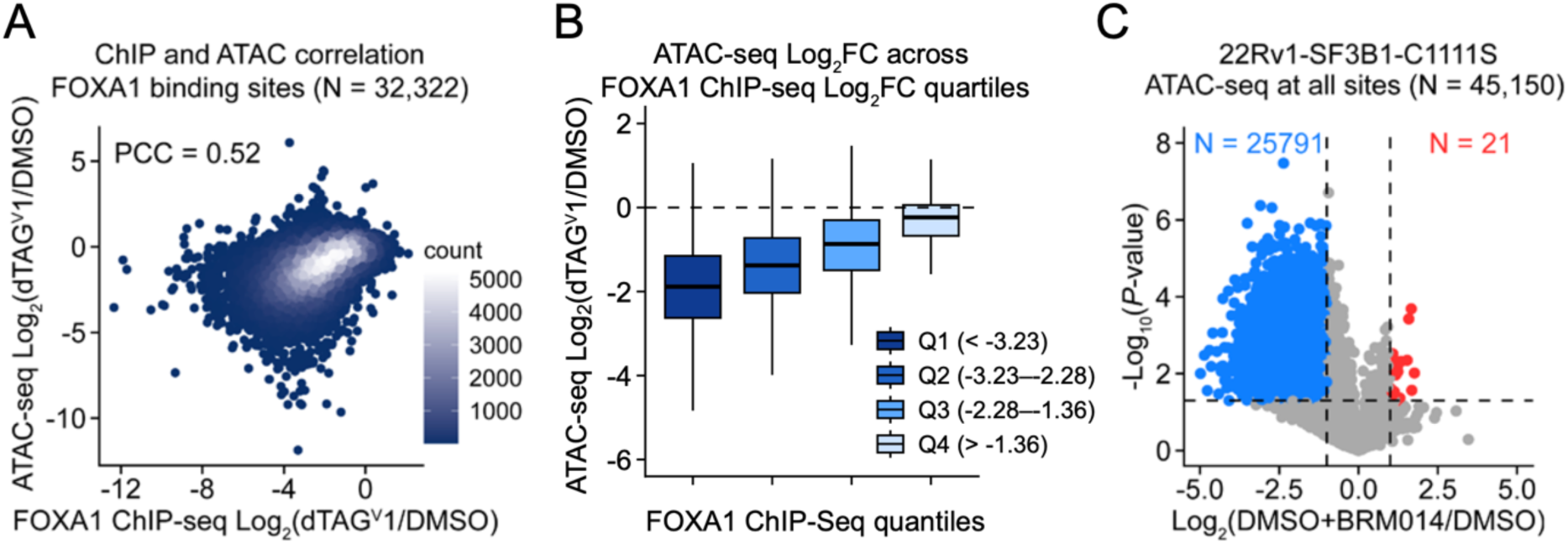
FOXA1 loss leads to chromatin accessibility decreases, related to. **Figure 2**. A. Scatter plot comparing dTAG-FOXA1 ChIP-seq and ATAC-seq changes (Log₂(dTAG^V^1/DMSO)) at all dTAG-FOXA1 binding sites (*N* = 32,322) after 1.5-h dTAG^V^1 treatment (500 nM). Pearson correlation coefficient (PCC) is shown. *n* = 3 biological replicates for ATAC-seq. B. Box plot showing ATAC-seq changes (dTAG^V^1/DMSO) stratified by FOXA1 ChIP-seq changes (quartiles of log₂(fold change) in FOXA1 ChIP-seq changes). Boxes indicate median and interquartile range, whiskers represent data range. C. Volcano plot of chromatin accessibility changes induced by 1 µM BRM014 for 4 h (1 µM BRM014 for 1 h, followed by DMSO for 3 h) compared to DMSO (DMSO for 1 h, followed by DMSO for 3 h) in 22Rv1-SF3B1-C1111S cells (*N* = 45,150 sites). Significantly changed sites (|log₂FC| > 1, *P* < 0.05) are shown in blue (decreased) or red (increased). *n* = 3 biological replicates for ATAC-seq.

**Figure S3:**
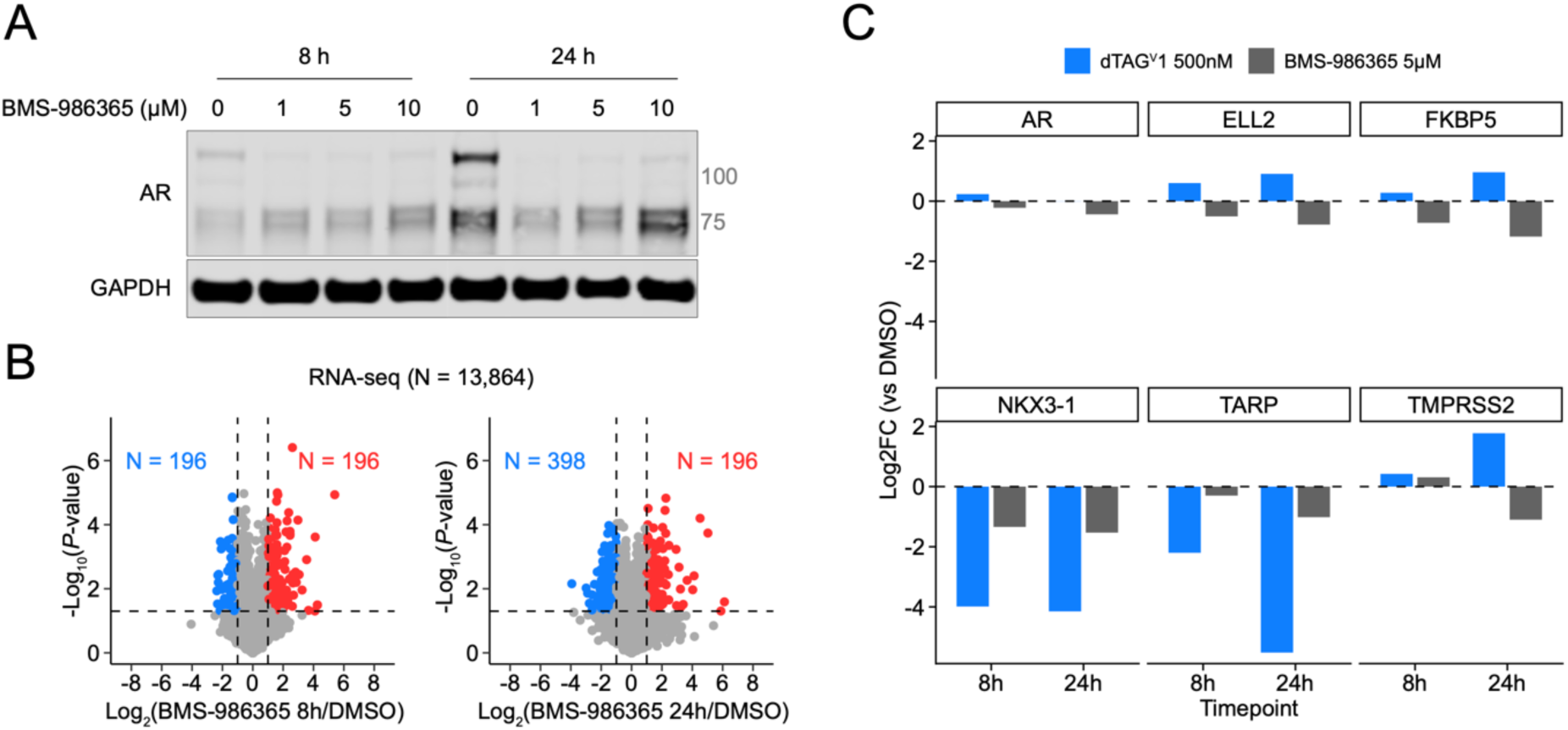
FOXA1 degradation causes bi-directional changes in transcription, related to. **Figure 3**. A. Immunoblot analysis of AR degradation. Immunoblot showing AR protein levels in 22Rv1-dTAG-FOXA1 cells treated with DMSO or indicated concentrations of BMS-986365 for 8 or 24 h. GAPDH used as a loading control. B. mRNA-seq data showing gene expression changes in 22Rv1-dTAG-FOXA1 cells treated with 5 µM BMS-986365 for 8 or 24 h (*N* = 13,864 transcripts). Transcripts significantly altered in expression following BMS-986365 treatment were defined as |log₂(BMS-986365/DMSO)| > 1 and *P* < 0.05, with red or blue for increased or decreased, respectively. *n* = 3 biological replicates. *P* values were determined by two-tailed Welch’s *t*-test. C. Bar plots of mRNA-seq log₂ fold changes (Treatment/DMSO) after 8 h or 24 h. Shown are *AR* and selected AR-associated targets (*ELL2, FKBP5, NKX3-1, TARP, TMPRSS2*).

**Figure S4.**
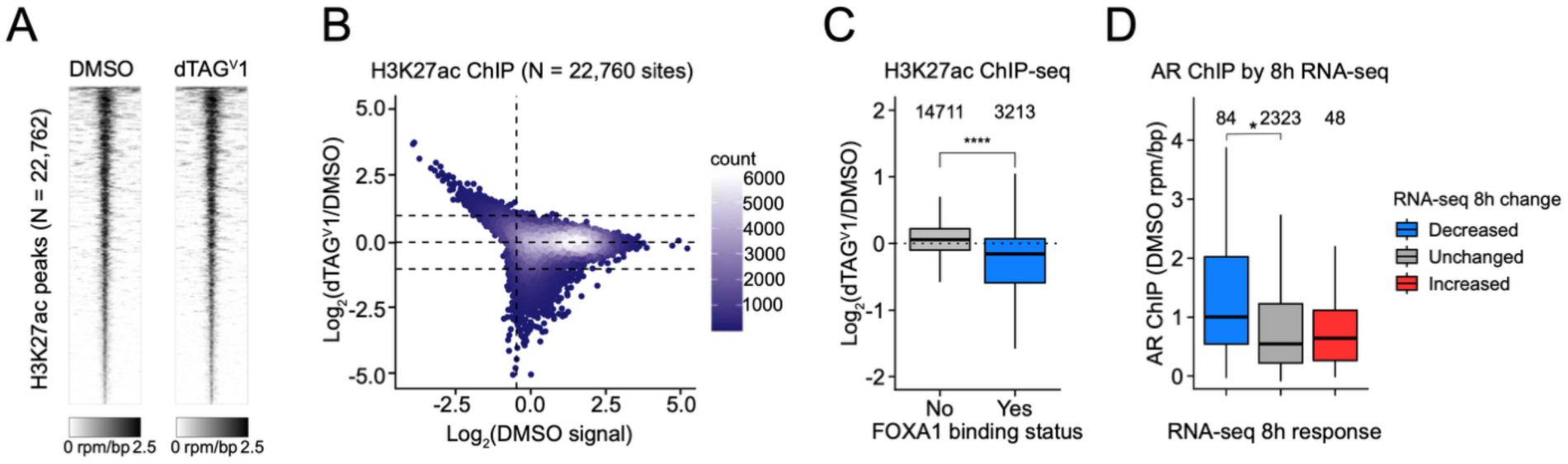
Context-dependent enhancer regulation by FOXA1, related to. **Figure 4**. A. DMSO rank-ordered heatmaps displaying H3K27ac ChIP-seq signal intensity across all H3K27ac sites in the presence or absence of 500 nM dTAG^V^1 treatment for 1.5 h in 22Rv1-dTAG-FOXA1. B. Mean-average (MA) plot showing log₂(dTAG^V^1/DMSO) changes in H3K27ac ChIP-seq signal as a function of DMSO signal (log_2_(rpm/bp)) in 22Rv1-dTAG-FOXA1 cells (*N* = 22,760 sites). Each point represents an H3K27ac peak, with color intensity indicating local point density. Dashed horizontal lines mark log₂ fold change thresholds of ±1 and 0, and the vertical line indicates the 10th percentile of DMSO signal. C. H3K27ac ChIP-seq changes at 1.5-h dTAG^V^1 treatment stratified by FOXA1 binding status (\*\*\*\**P* < 0.0001, Welch’s *t*-test). Sites with > 1 rpm/bp in at least one H3K27ac ChIP-seq sample were included in analysis (*N* = 17,924 sites). D. Baseline signal (rpm/bp in DMSO) for AR ChIP-seq in 22Rv1 parental cells (data from GEO, GSE275277) at FOXA1 regulatory elements, grouped by in 22Rv1-dTAG-FOXA1 RNA-seq-defined target gene response at 8 h (as in Figure 4C-F). Gene response category thresholds are defined in Figure 4A. (\**P* < 0.05, \*\**P* < 0.01, \*\*\**P* < 0.001, \*\*\*\**P* < 0.0001, Welch’s *t*-test).

**Table S1:**
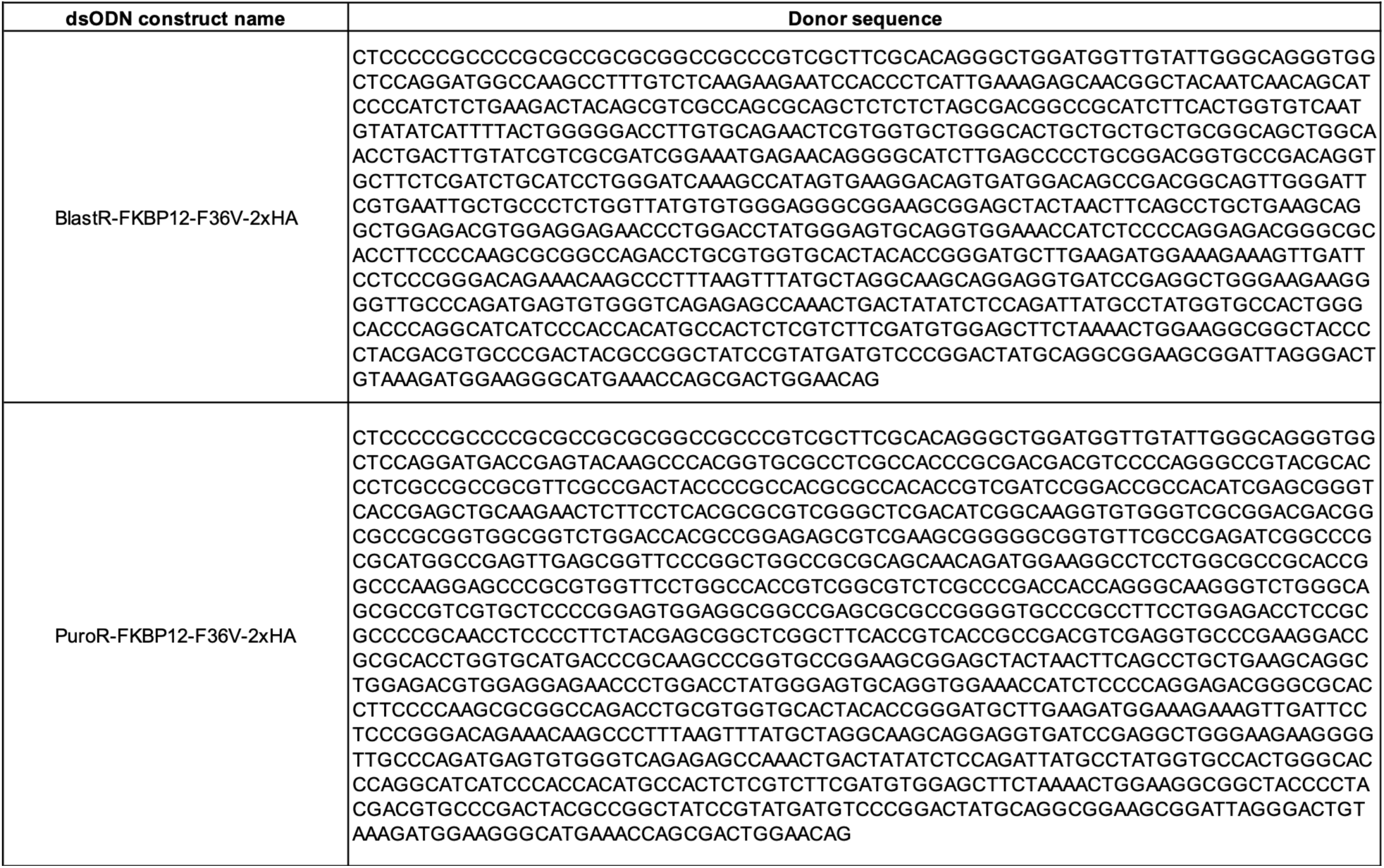
FKBP12-F36V-2xHA dsODN sequences.

