## Supplementary material for "Dissecting FOXA1 pioneering function by acute pharmacological degradation": Materials and Methods

#### Experimental models and subject details

#### Cell lines and culture conditions

22Rv1 prostate cancer cells (ATCC), 22Rv1-Cas9 cells, and 22Rv1-SF3B1-C1111S were maintained in RPMI-1640 (with L-glutamine, Gibco) supplemented with 10% fetal bovine serum (FBS) and 1X Antibiotic-Antimycotic (100X stock, Gibco) at 37°C with 5% $CO_2$. LentiX 293T cells were maintained in DMEM (with L-glutamine, Gibco) supplemented with 10% fetal bovine serum (FBS) and 1X Antibiotic-Antimycotic (100X stock, Gibco) at 37°C with 5% $CO_2$. Cells were regularly tested for mycoplasma contamination (Lonza MycoAlert). For all experiments, cells were seeded to ensure logarithmic growth during treatments.

#### CRISPR-mediated generation of 22Rv1-dTAG-FOXA1 cells

CRISPR/Cas9-mediated genome editing was performed using a Neon Transfection System (Thermo Fisher Scientific) according to the manufacturer's guidelines. Single guide RNAs (sgRNAs) targeting the N-terminus of *FOXA1* locus (custom made from IDT; 5'-CUUCACAGUUCCUAACAUCC-3') and double-stranded DNA homology donor oligonucleotides (dsODNs) were obtained from IDT and resuspended to 100 µM and 1 µg/µL, respectively. The double-stranded oligodeoxynucleotide (dsODN) donor contained the sequence encoding either the puromycin or blasticidin resistance cassette, P2A self-cleaving peptide, the $FKBP12^{F36V}$ degron, and 2×HA epitope tag, flanked by ~50bp homology arms targeting the N-terminus of the *FOXA1* locus (full donor sequences listed in Table S1). For ribonucleoprotein (RNP) complex formation, 2.9 µL of Alt-R™ S.p. HiFi Cas9 Nuclease V3 (61 µM; IDT), 1.3 µL duplex buffer (Thermo Fisher Scientific), and 3.5 µL sgRNA (100 µM) were combined (2:1 sgRNA:Cas9 molar ratio) and incubated at 37°C for 15 minutes to form a 20 µM RNP complex. Meanwhile, 200 k 22Rv1 cells were harvested, washed once in PBS to remove antibiotics, and resuspended in 10 µL of Buffer R per electroporation. Each electroporation reaction contained 1 µL of the pre-formed RNP (20 pmol, ~2 µM final) and 1 µL of each dsODN donor (1 µg/µL puromycin and blasticidin resistance cassettes). The RNP and dsODN mixture was gently combined with the cell suspension and aspirated into a Neon pipette tip. Electroporation was performed using the following parameters: 1250 V, 20 ms pulse width, and 2 pulses. Immediately after electroporation, cells were transferred into antibiotic-free RPMI-1640 supplemented with 10% FBS and incubated at 37°C. After 24 hours, 500 µL of media containing anti-anti was added to each well. Selection with puromycin (1:5000) was initiated once cells reached approximately 80% confluency. After one week of selection, blasticidin (1:1000) was added for an additional week to select for double-resistant clones. Genomic integration was confirmed by PCR, nanopore sequencing (premium PCR, Plasmidasaurus) and immunoblot analysis of FOXA1, and successfully edited populations were used directly for degradation assays.

**Generation of 22Rv1-Cas9 cells**

22Rv1-Cas9 cells were generated by lentiviral transduction with pFUGW Cas9-BFP[1] (a gift from Xiaowei Zhuang; Addgene plasmid # 127396 ; http://n2t.net/addgene:127396 ; RRID:Addgene_127396). Cells were transduced with Cas9-BFP virus via spinfection at 1000 x g for 1 h at room temperature with the addition of 8μg/mL polybrene reagent (MilliporeSigma). Following transduction, cells were expanded, analyzed by flow cytometry, and sorted into low, medium, and high BFP-expressing populations. Each population was expanded separately, and expression levels were validated by western blot analysis. The population showing optimal Cas9 expression was selected for subsequent genome editing experiments.

**Generation of 22Rv1-SF3B1-C1111S cells**

22Rv1-SF3B1-C1111S cells were generated by transfecting cells with a pre-assembled ribonucleoprotein (RNP) complex and single-stranded oligodeoxynucleotide (ssODN) donor template designed for homology-directed repair (HDR), as previously described[2].

### Methods

**Cancer Dependency Map (DepMap) analyses**

Cancer dependency and gene expression data were obtained from the DepMap portal (https://depmap.org). Gene dependency scores from genome-scale CRISPR-Cas9 loss-of-function screens (CRISPRGeneEffect; Chronos) and RNA-seq expression values (TPM; public 24Q4 release) were used for analyses shown in Figures 1A and B. Cell lines were grouped by lineage based on DepMap annotations. For lineage-specific analyses, Chronos gene effect scores were compared between (i) prostate cancer cell lines versus all other lineages and (ii) breast cancer cell lines versus all other lineages on a per-gene basis. For each gene, the differential gene effect was calculated as the difference in mean gene effect scores between the lineage group and all other lineages (mean_lineage − mean_other), and statistical significance was assessed using a two-sided Welch's t-test (unequal variance). *P* values were plotted as -log10(*P*), with FOXA1 highlighted. For cross-lineage analyses, FOXA1 gene effect scores were plotted against FOXA1 expression values across all available cell lines and colored by lineage.

**Small-molecule treatments**

The following compounds were used

- dTAG$^{V}$1 (VHL-based degrader of FKBP12$^{F36V}$; Tocris)
- BRM014 (SMARCA2/4 ATPase inhibitor; Sigma)
- WX-02-23 (tryptoline acrylamide FOXA1 ligand)
- BMS-986365 (dual AR degrader/antagonist; MedChemExpress)

WX-02-23 was synthesized by WuXi AppTec and synthesis and characterization have been reported in previous studies[3].

**Lentivirus production**

Lentiviral production was performed in Lenti-X™ 293T cells via co-transfection with packaging plasmids pMD2.G and psPAX2 (Addgene), polyethylamine (PEI) transfection agent, and the lentiviral expression plasmid of interest (5 μg). Lenti-X cells were plated in anti-anti free DMEM 10% FBS media and then switched to media containing 1X anti-anti 24 h following transfection. Virus was collected at 48 and 72 h post transfection filtered through a 0.45 μM membrane and concentrated overnight with Lenti-X concentrator (Takara).

**CRISPR-Cas9 competitive growth assays**

sgRNA sequences were cloned into the LRG backbone[4] (a gift from Christopher Vakoc, Addgene plasmid #65656; http://n2t.net/addgene:65656; RRID:Addgene_65656) for co-expression of an sgRNA and GFP. Virus was produced in Lenti-X cells as detailed above. 22Rv1-Cas9-expressing cells were transduced in triplicate with the corresponding sgRNA-LRG virus at 30-60% efficiency in 96-well plates. Transduced cells were passaged and subjected to flow cytometry to measure GFP-positive percentage every 3 to 4 days. sgRNA sequences, which were taken from the Brunello sgRNA library[5] are as follows: AAVS1 5'-GGGGCCACTAGGGACAGGAT-3'; RPS19 5'-GTAGAACCAGTTCTCATCGT-3'; Luciferase 5'-CCCGGCGCCATTCTATCCGC-3'; FOXA1-sg2 5'-GTCCGGGTGCAGCGTCCAGT-3'.

**Western Blotting**

Whole-cell lysates were prepared in RIPA buffer (Thermo Scientific) supplemented with 1X Halt™ Protease Inhibitor Cocktail, EDTA-Free (Thermo Scientific) and 1X benzonase nuclease (Sigma-Aldrich). Protein concentrations were determined using the Pierce™ BCA Protein Assay Kit, and samples were normalized to load 30μg protein per well per sample. Protein samples were resolved on Bolt™ Bis-Tris Plus 4–12% Mini Protein Gels (Thermo Fisher Scientific, 1.0 mm, WedgeWell™ format) and transferred with 1X Bolt™ Transfer Buffer (Thermo Fisher Scientific) with 10% methanol to nitrocellulose membranes (Thermo Scientific, 0.45 μm). Membranes were blocked in 5% nonfat milk in TBST (1X TBS, 0.1% Tween) for 1 hour and incubated overnight at 4 °C with primary antibodies against FOXA1 (1:500, Cell Signaling Technology), HA (1:1000, Cell Signaling Technology), and GAPDH (1:5000, Sigma-Aldrich). Blots were washed 3 x 10 min in TBST. After washing, blots were incubated with IRDye 680RD Goat Anti-Mouse IgG and IRDye 800CW Goat Anti-Rabbit IgG secondary antibodies (1:5000; LI-COR) for 1 h at room temperature. Blots were washed again 3 x 10 min in TBST. Blots were visualized using a LI-COR Odyssey CLx imaging system. Representative blots are shown in Figures 1E, F, and S1C.

**Protein-directed activity-based protein profiling**

Protein-directed ABPP was performed as previously described[6]. In short, 22Rv1-dTAG-FOXA1 cells were treated with DMSO or dTAG$^{V}$1 (500nM; 0.5 or 2 h), followed by the addition of alkyne probes (CJR-6A or CJR-6B; 5 μM; 1 h). Cells were washed with ice-cold DPBS (3x) and collected. Cell pellets were resuspended in DPBS and lysed by probe-sonication (2 x 15 pulses; 10% power output). Proteome was normalized to 2 mg/ml in 500 μL, and the alkyne probe labeled proteins were conjugated to biotin-PEG4-azide via Cu(I)-catalyzed azide–alkyne cycloaddition (CuAAC; 2 mM CuSO4, 100 μM Biotin-PEG4-azide, 1 mM tris(2-carboxyethyl)phosphine (TCEP), 100 μM tris((1-benzyl-4-triazolyl)methyl)amine (TBTA); 1 h). After chloroform/methanol precipitation and centrifugation (16,000 g, 10 min, 4 °C), solvents were aspirated and the protein-disk was washed with 1 mL ice-cold MeOH and resuspended in 0.5 mL resuspension buffer (8 M urea, 0.2 % SDS in DPBS). Peptides were reduced (10 mM DTT; 15 min; 65 °C) and alkylated (20 mM iodoacetamide; 30 min; 37 °C). Next, 130 μL 10% SDS was added, samples were diluted to 6 mL in DPBS and incubated with 100 μL of streptavidin-agarose bead slurry for 90 minutes. Beads were washed (2 x 0.2 % SDS in DPBS, 1 x DPBS, 2 x $H_2O$, 1 x 200 mM EPPS, pH 8.0) and enriched proteins were on-bead digested with mass-spectrometry grade trypsin in buffer (2 M urea in 200 mM EPPS, 1 mM $CaCl_2$) overnight at 37 °C. Digested peptides were TMT-labeled (16-plex) for 90 minutes and quenched by hydroxylamine. The TMT-labeled peptides were desalted by Sep-Pak C18 cartridges (Waters) and high-pH fractionated using peptide desalting spin columns (Thermo). A total of 10 fractions were analyzed by Orbitrap Eclipse Tribid Mass Spectrometer with Xcalibur v.4.3 (see below for LC-MS instrumentation and analysis).

**TMT liquid chromatography-mass-spectrometry (LC-MS) analysis**

TMT LC-MS analyses were performed as previously described using the same instruments and workflow settings[3,6]. All samples were analyzed by Orbitrap Eclipse Tribrid Mass Spectrometer coupled to an UltiMate 3000 Series Rapid Separation LC system and autosampler (Thermo Scientific Dionex). Data was acquired using an MS3-based TMT method (see ref. Lazear et al.[3] for the detailed settings) and the RAW files were uploaded and processed by Integrated Proteomics Pipeline (IP2, v.6.7.1), and searched using the ProLuCID algorithm95 using a reverse concatenated, non-redundant variant of the Human UniProt database (release 2016). Cysteine residues were searched with a static modification for carboxyamidomethylation (+57.02146 Da). N-term and lysine static modification from the tags (+304.2071 Da for 16-plex) were included. Peptides were required to be at least 6 amino acids long and filtered to through DTASelect (version 2.0) to obtain a peptide false-positive rate below 1%. The MS3-based peptide quantification was performed with reporter ion mass tolerance set to 20 ppm with Integrated Proteomics Pipeline (IP2).

**Data processing of protein-directed ABPP**

The census output files from IP2 were further processed to calculate enrichment engagement ratios (probe vs probe) by dividing each TMT reporter ion intensity by the sum of intensity for all the channels. Each spectrum-peptide match was grouped based on protein ID, excluding peptides with summed reporter ion intensities < 10,000, coefficient of variation of > 0.5, < 2 unique peptides per protein ID. TMT reporter ion intensities were normalized to the median summed signal intensity across channels.

**ChIP-seq**

22Rv1-dTAG-FOXA1 or parental 22Rv1 cells were treated with DMSO or 500 nM dTAG$^{V}$1 for 1.5 hours (15-cm dish; ~70-80 % confluency; 30 million cells). Cells were washed with PBS, trypsinized, and crosslinked with 1.1 % formaldehyde (final) in 50 mM HEPES pH 7.5, 100 mM NaCl, 1 mM EDTA, and 0.5 mM EGTA for 10 minutes at room temperature. Crosslinking was quenched with 125 mM glycine for 5 minutes. Cells were washed three times with ice-cold PBS and snap-frozen at –80 °C until processing. For chromatin preparation, pellets were resuspended in Lysis Buffer 1 (LB1; 50 mM HEPES pH 7.5, 140 mM NaCl, 1 mM EDTA, 10 % glycerol, 0.5 % NP-40, 0.25 % Triton X-100, 1× Halt™ Protease Inhibitor Cocktail (Thermo Fisher)) and rotated for 10 minutes at 4 °C, followed by centrifugation at 1,400 × g for 5 minutes at 4 °C. Pellets were then washed with Lysis Buffer 2 (LB2; 10 mM Tris pH 7.5, 200 mM NaCl, 1 mM EDTA, 0.5 mM

EGTA, 1× protease inhibitor) and resuspended in Sonication Buffer (50 mM HEPES pH 7.5, 140 mM NaCl, 1 mM EDTA, 1 mM EGTA, 1 % Triton X-100, 0.1 % sodium deoxycholate, 0.5 % SDS, 1× protease inhibitor). Chromatin was sheared using a Diagenode Bioruptor (25 cycles, 30 sec on/off, 10 °C). 50μL of whole cell extract (input) for the DMSO sample was moved to a new tube and stored at -20 °C until the reverse crosslinking step. Sheared chromatin was diluted to 0.1 % SDS and incubated overnight at 4 °C with antibodies conjugated to Dynabeads™ Protein G (Thermo Fisher): 10 μg FOXA1 (Novus Biologicals) or 5 μg H3K27ac (Diagenode). Beads were washed twice with Sonication Buffer (without SDS), once with High-Salt Sonication Buffer (500 mM NaCl, without SDS), once with LiCl Wash Buffer (20 mM Tris pH 7.5, 1 mM EDTA, 250 mM LiCl, 0.5 % NP-40, 0.5 % sodium deoxycholate), and once with TE + 50 mM NaCl. Bound chromatin was eluted in 200μL Elution Buffer (50 mM Tris pH 7.5, 10 mM EDTA, 1 % SDS) at 65 °C for 15 minutes. The input sample was thawed and 150μL of elution buffer was added before crosslinking with the other samples at 65 °C overnight (13 h). Samples were treated sequentially with 8μL 10mg/mL RNase A (37 °C, 2 h), 7μL of 300mM CaCl2 in 10mM Tris at pH 8, and 4μL 20mg/mL Proteinase K (55 °C, 30 min), followed by extraction with phenol:chloroform:isoamyl alcohol (25:24:1). The aqueous phase was isolated and DNA precipitated at -80 °C for 2-3 h. DNA fragments (300-500 bp) were purified and processed for library preparation using the ThruPLEX® DNA-seq Kit (Takara Bio), indexed with DNA Unique Dual Index Kit -24U Set A (Takara), cleaned with AMPure XP beads (Beckman Coulter), and sequenced on an Illumina NextSeq 2000 (single-end, 75 bp).

**ChIP-seq analysis**

Sequencing reads were aligned to human genome build hg19 and RefSeq genes by Bowtie2[7] with the following parameters: -N 1 -p 10 -x. Using the unenriched input sample as background, the signal-enriched regions were identified by Model-based Analysis of ChIP-seq (MACS)[8] peak-finding algorithm (v1.4.1) with the *P* value threshold of 1e-9. For FOXA1 ChIP samples, a merged peak list including all FOXA1-binding sites from all relevant conditions (DMSO, dTAG$^{V}$1 for 22Rv1-dTAG-FOXA1 cells, and DMSO for 22Rv1 parental cells) was created with bedtools (v2.17.0) for downstream analyses (Figure 1F, S1E). A merged peak list including all dTAG-FOXA1-binding sites from all relevant conditions (DMSO, dTAG$^{V}$1 for 22Rv1-dTAG-FOXA1 cells) was created with bedtools (v2.17.0)[9] for other dTAG-FOXA1 specific downstream analyses (Figure S1F). For the H3K27ac ChIP samples, a merged peak list including all H3K27ac-binding sites from all relevant conditions (DMSO and dTAG$^{V}$1) was created with bedtools (v2.17.0) for downstream analyses. Using the read density calculator Bamliquidator (v1.0)[10] (http://github.com/BradnerLab/pipeline/wiki/bamliquidator) to map sequencing reads to designated loci with a 200 bp extension window in either direction differential analyses were performed across samples. Samples were normalized by total number of mapped reads (reads per million, rpm) and read density per base pair was calculated (rpm/bp). All box plots represent 25-75 percentile with whiskers extending 1.5 interquartile range (IQR) and the center line represents the median. For calculation of average FOXA1 peak intensity (Figure S1F), DMSO- and dTAG$^{V}$1-treated dTAG-FOXA1 ChIP-seq samples were mapped to a combined peak set derived from both conditions. Signal was aggregated in ±5 kb windows centered on each peak and divided into 200 bins. For each sample, signal was averaged across all peaks at each bin and plotted as the average peak intensity profile.

Publicly available AR ChIP-seq data were obtained from the Gene Expression Omnibus (GEO; GSE275777). Raw sequencing reads were aligned to the human genome (hg19) using Bowtie2 with the same parameters described above. For downstream analyses, aligned AR ChIP-seq BAM files were mapped to FOXA1-defined regulatory elements (defined above) using Bamliquidator with a 200 bp extension window on either side of each site. AR ChIP-seq data were used exclusively to assess AR occupancy these FOXA1 sites and were not subjected to AR peak calling or genome-wide differential analysis. Results are shown in Figure S4D.

**Cell proliferation assays**

Cells were seeded at 1,000 cells per well in 40 µL of growth medium in 384-well clear-bottom, white-wall plates (Corning). 40 nL of DMSO or compounds (prepared at 1,000× final concentration) were transferred using an Echo 650 acoustic dispenser from 384-well Low Volume Echo-qualified source plates (Beckman Coulter, Cat. #P-05525). A baseline plate was read on day 0 for normalization purposes. Plates were incubated for 120 h at 37 °C. Cell viability was measured using the CellTiter-Glo Luminescent Cell Viability Assay (Promega) according to the manufacturer’s instructions. Luminescence was recorded on a CLARIOstar plate reader (BMG Labtech). Average day 0 RLU values were subtracted from day 5 RLU, and data were normalized to DMSO-treated controls at day 5.

**Cell growth assays**

Cells were seeded in triplicate and treated as indicated. Viable cell counts were determined using a Countess II automated cell counter (Thermo Fisher Scientific) at daily intervals for 5 days. Cell numbers were normalized to day 0. Mean ± SD values are shown from three biological replicates (Figures S1F).

**ATAC-seq**

ATAC-seq was performed using the Active Motif ATAC-Seq Buffer Set and 96-well Pre-Indexed Assembled Tn5 Transposomes according to the manufacturer's protocol, with minor modifications. 22Rv1-dTAG-FOXA1 cells were treated with DMSO or 500 nM dTAG$^{V}$1 for 1.5 hours (24-well plates, 70–80 % confluency). 22Rv1-SF3B1-C1111S cells were pre-treated with 1 µM BRM014 for 1 h, followed by 20 µM WX-02-23 for 3 h (24-well plates, 70–80 % confluency). After treatment, cells were washed twice with ice-cold PBS and lysed directly in-plate with 150 µL ATAC lysis buffer (Active Motif). Lysates were centrifuged at 500 × g for 10 min at 4 °C, and the supernatant was carefully removed to isolate nuclei. Each nuclei pellet was resuspended in 46 µL Tagmentation Master Mix (1× Tagmentation Buffer, 1× PBS, 0.01 % digitonin, 0.1 % Tween-20). For indexing, each sample was combined with a distinct well of the Pre-Indexed Assembled Tn5 Transposome plate, ensuring unique barcode assignment per sample. Plates were sealed, gently vortexed, quick-spun, and incubated at 37 °C for 30 min with gentle mixing every 10 min. DNA was purified and libraries were prepared according to the manufacturer's protocol. Libraries were cleaned by SPRI-bead purification and gel extraction to enrich for fragments < 1 kb, then multiplexed and sequenced on an Illumina NextSeq 2000 using 50-bp paired-end reads.

**ATAC-seq analysis**

Sequencing reads were aligned to the human reference genome (hg19) and RefSeq gene annotations using Bowtie2 (v2.5.1)[7] with the following parameters: bowtie2 --local --very-sensitive-local --no-mixed --no-discordant --dovetail --phred33 -I 10 -X 700 --threads 12 -x. Unmapped, duplicate, and low-quality reads were removed using Sambamba (v0.7.1)[11]. Reads mapping to the mitochondrial genome or ENCODE blacklist regions (*wgEncodeDacMapabilityConsensusExcludable.bed.gz*) were excluded using SAMtools (v1.10)[12]. Peak calling was performed with MACS (v2.1.0) using the parameters: macs2 callpeak -q 0.01 --nomodel --shift -100 --extsize 150. Adjacent peaks within 1 kb were merged to define consolidated accessible regions. A union peak set was generated by merging all ATAC-seq peaks detected across samples for downstream analyses. Differential analyses were performed with the same method used for ChIP-seq experiments as described in the previous section. To minimize noise and restrict analyses to high-confidence sites, regions with an average ATAC signal of at least 1 rpm/bp in any treatment condition (based on biological triplicates) were retained. Differential accessibility was calculated using $\log_2$ fold-change ($\log_2$FC) and *P*-value thresholds, with significantly changed sites defined as $\log_2$FC > 1 or < -1 and $P < 0.05$. All box plots represent 25-75 percentile with whiskers extending 1.5 interquartile range (IQR) and the center line represents the median.

**3' mRNA-seq**

22Rv1-dTAG-FOXA1 cells (70 % confluency; 6-well plates) were treated with the indicated compounds for the specified durations. Total RNA was extracted using the RNeasy Plus Mini Kit (Qiagen) with 0.1 % (v/v) β-mercaptoethanol (BME) added to all buffers. RNA concentration was quantified using the Qubit Broad Range RNA Assay Kit (Thermo Fisher Scientific). For library preparation, 500 ng of total RNA per sample was processed using the QuantSeq 3′ mRNA-seq V2 Library Prep Kit FWD with UDI (Lexogen) according to the manufacturer's instructions. Samples were purified using the magnetic bead purification module (Lexogen). Libraries were multiplexed and sequenced on an Illumina NextSeq 2000 platform to generate 100 bp single-end reads.

**3' mRNA-seq analysis**

Raw sequencing data (FASTQ format) were processed using the SLAMdunk[13] pipeline for alignment to the human genome (hg19) and counts-per-million (CPM) quantification. Transcripts with substantial expression (average >3 CPM in at least one treatment group) were retained for downstream analysis. All downstream analyses were performed in R, and code is available upon request.

**Integrative multi-omic analysis**

FOXA1 ChIP-seq, ATAC-seq, and H3K27ac datasets were integrated using GenomicRanges[14] and custom R scripts. FOXA1 regulatory elements were identified using DMSO dTAG-FOXA1 ChIP-seq peaks and peaks were stitched using ROSE2[15,16]. Regulatory elements were provisionally associated with the nearest annotated transcription start site based on linear genomic distance. BAM files from dTAG-FOXA1 ChIP-seq, ATAC-seq, and H3K27ac datasets in 22Rv1-dTAG-FOXA1 cells were mapped to the set of FOXA1-defined regulatory elements using the methods used for ChIP-seq explained above. Aligned AR ChIP-seq BAM files were mapped to FOXA1-defined regulatory elements solely to assess AR occupancy at these sites (Figure S4D). When integrating mRNA-seq data with enhancer signal, the transcript with the highest average DMSO 8 h baseline CPM was selected to represent each gene. In cases where multiple enhancers were assigned to the same gene, the enhancer with the highest DMSO dTAG-FOXA1 ChIP signal was selected. $Log_2$ fold-change ($Log_2FC$) and RPM/BP values were used to compute correlations, stratify regulatory elements by target gene transcriptional response, and generate box plots (Figures 4A–F, S4D). Genome browser tracks were generated from hg19-aligned BAM files using a custom Python-based plotting workflow. Statistical significance was determined using Welch's *t*-test unless otherwise stated. Pearson correlation coefficients were calculated for pairwise comparisons. All plots were generated in R (v4.4.1).

**Data availability**

All raw sequencing data have been deposited in GEO under accession number GSE315959. The mass spectrometry proteomics data have been deposited to the ProteomeXchange Consortium via the PRIDE[17] partner repository with the dataset identifier PXD074757. Processed proteomics and sequencing data are provided in Supplementary Dataset S1 and S2, respectively. Custom analysis scripts and plotting code are available upon request.
